# Polycomb response elements reduce leaky expression of Cas9 under temperature-inducible *Hsp70Bb* promoter in *Drosophila melanogaster*

**DOI:** 10.1101/2022.08.29.505746

**Authors:** Natalie Warsinger-Pepe, Carly Chang, Connor R. Desroberts, Omar S. Akbari

**Affiliations:** School of Biological Sciences, Department of Cell and Developmental Biology, University of California, San Diego, La Jolla, CA 92093; School of Biological Sciences, University of California, San Diego, La Jolla, CA 92093

**Keywords:** inducible, Cas9, heat-shock, PREs, Polycomb response elements, Drosophila melanogaster

## Abstract

Heat shock inducible expression of genes through the use of heat inducible promoters is commonly used in research despite leaky expression of downstream genes of interest without targeted induction (i.e. heat shock). The development of non-leaky inducible expression systems are of broad interest for both basic and applied studies, to precisely control gene expression. Here we characterize the use of Polycomb response elements and the inducible *Heat shock protein 70Bb* promoter, previously described as a non-leaky inducible system, to regulate Cas9 endonuclease levels and function in *Drosophila melanogaster* after varying both heat shock durations and rearing temperatures. We show that Polycomb response elements can significantly reduce expression of *Cas9* under *Heat shock protein 70Bb* promoter control using a range of conditions, corroborating previously published results. We further demonstrate that this low transcript level of heat-induced *Cas9* is sufficient to induce mutant mosaic phenotypes. Incomplete suppression of an inducible *Cas9* system by Polycomb response elements with no heat shock suggests that further regulatory elements are required to precisely control *Cas9* expression and abundance.

## Introduction

Temporal control of expression through inducible gene expression systems (IGES) is a useful genetic strategy to understand gene function. One commonly utilized IGES is the heat-shock response system, whereby a heat shock promoter is placed upstream of a transgene, or gene of interest, which can then be activated by endogenous heat shock proteins when the environment temperature shifts from lower temperatures to 37°C (heat shock) (Pelham 1982; Lindquist 1984). Despite the power and utilization of this tool, the heat shock promoter exhibits leaky expression, where promoter activity is seen at temperatures below 37°C (Lindquist 1984; Curtin *et al*. 2008; Naidoo and Young 2012; Fujita *et al*. 2014; Sloan and Barres 2014; Akmammedov *et al*. 2017; Nandy *et al*. 2019). The *Drosophila Heat shock protein 70Bb* (*Hsp70Bb*) promoter is regularly used for temporal control of gene expression through heat shock in many organisms despite its leakiness (Corces *et al*. 1981; Pelham and Bienz 1982; Pelham 1982; Spena *et al*. 1985; Wei *et al*. 1986; Kust *et al*. 2014). Akmammedov and colleagues utilized *bithoraxoid* (*bxd*) Polycomb response elements (PREs) (Simon *et al*. 1993), which silence adjacent genes through Polycomb group (PcG) protein deposition of H3K27me3 silencing marks (Beisel and Paro 2011), upstream the *Hsp70Bb* promoter to significantly suppress leaky expression of a downstream gene, *lacZ* (Akmammedov et al. 2017). Despite this significant reduction in leaky expression, PRE repeats appear to not be widely used.

Suppressing leaky expression of genes under *Hsp70Bb* promoter control has the potential to aid in basic biological studies of pathway mechanisms as well as applied systems for pest control. Temporal and spatial control of gene expression can aid in understanding details of gene and/or pathway function by regulating timing and/or dosage of gene transcription and products. Temporal control of sterile insect technologies like precision-guided sterile insect technique (pgSIT) can be a powerful and financially impactful tool for scaling up large populations of pest species for release and population control (Kandul *et al*. 2019, 2021, 2022). Heat-shock inducible *Cas9* has been used in multiple organisms to induce programmed mutations, including in rice (Nandy *et al*. 2019) and *Drosophila* (Kandul et al. 2021), however these systems still reveal *Cas9* activity without heat shock. In this study, we characterize the effectiveness of PREs upstream *Hsp70BbCas9* in suppressing leaky *Cas9* expression without heat shock. We characterize the ability of *Hsp70BbCas9* to induce mutations in the presence of guide RNAs (gRNAs) against three separate gene targets (*w, ey, Ser*) after varying heat shock durations (0, 30 minutes, 1 hour, 2 hours), and at three different rearing temperatures (18°C, 21°C, 26°C). Through this genetic characterization, we show that PREs are able to drastically reduce *Cas9* expression under the majority of conditions tested, yet low levels of functional Cas9 are often sufficient to induce mutant mosaic phenotypes.

## Materials and Methods

### Fly husbandry and strains

*D. melanogaster* stocks were either reared in 18°C, 21°C, or 26°C incubators with standard light/dark cycles on Texas media from the UCSD *Drosophila* Recharge facility (Fly Kitchen). Stocks were reared at their subsequent temperature for at least 3 generations before being utilized in experiments. The following fly stocks from the Bloomington *Drosophila* Stock Center were used: p{TKO.GS02468}attP40/II “ *sgRNA: w*” (BDSC 79543), p{WKO.1-G12}attP40/II “ *sgRNA: ey*” (BDSC 82495), M{WKO.p1-B12}ZH-86Fb/TM3,Sb^1^ “ *sgRNA: Ser*” (BDSC 84169), *w*^*1118*^; PBac{y^+mDint2^=vas-Cas9}VK00027 “ *vasaCas9*” (BDSC 51324), y[1] M{w^+mC^=nos-Cas9.P}ZH-2A w* “ *nosCas9*” (BDSC 54591). P{y[+t7.7]=CaryP}attP2. *Hsp70BbCas9* was generated by Nikolay Kandul and Junru Liu (BDSC 92793). PRE-Hsp70Bb-Cas9 plasmid was generated as described below. Embryo injections to generate both *Hsp70BbCas9* and *PRE-Hsp70BbCas9* transgenic animals were carried out at Rainbow Transgenic Flies, Inc. (http://www.rainbowgene.com) using ϕC31-mediated integration into the same genomic attP2 location (BDSC # 8622) and balanced using standard balancer chromosomes. All stocks are listed in the Reagents Table.

### Transgene construction

For construction of PRE-Hsp70Bb-Cas9, Hsp70Bb-Cas9-T2A-eGFP (Kandul *et al*. 2021) was PCR amplified with NEB Q5 High-Fidelity 2X Master Mix (M0492) using primers 1157_onestep_p1 and 1157_onestep_p2. PRExpress (Akmammedov *et al*. 2017) was obtained from Addgene (122486). The PRE repeats of PRExpress were removed using Kpn-I and Nhe-I and purified using Zymoclean Gel DNA Recovery Kit (Genesee Scientific #11-301). This fragment was subcloned with the above PCR fragment using Gibson enzymatic assembly (Gibson *et al*. 2009) to generate PRE-Hsp70BbCas9_1.0 (Addgene 190795). Gypsy insulator elements were subsequently cloned into PRE-Hsp70BbCas9_1.0 through two Gibson cloning events to generate PRE-Hsp70BbCas9_1.2 (Addgene 190796) with one gypsy insulator element (KpnI digest, PCR with 1157A_onestep_for and 1157A_onestep_rev) and subsequently PRE-Hsp70BbCas9_1.3 (Addgene 190797) with two gypsy insulator elements (AfeI digest, PCR with 1157B_onestep_for and 1157B_onestep_rev). This final product, PRE-Hsp70BbCas9_1.3 (referred to as PRE-Hsp70Bb-Cas9), was used for transgenesis. Plasmid sequences were verified through Sanger sequencing to ensure the only differences between Hsp70Bb-Cas9-T2A-eGFP and PRE-Hsp70BbCas9_1.3 were the presence of the PREs and gypsy insulator elements (Figure S1). All Sanger sequencing was performed by either Retrogen, Inc. or GENEWIZ, Inc. All primers and plasmids are listed in the Reagents Table. At every cloning step, the PRE repeat size was verified (∼2.5kb) through restriction digestion with Kpn-I and Xho-I, as these repeats are often unstable.

### Mutant generation with cell-specific promoters

To observe mutant phenotypes from the chosen gRNA strains, gRNA stocks were crossed to either *nanos-Cas9* (*nosCas9*) or *vasaCas9* stocks, and the rate of mutant phenotype generation was assessed. All crosses were performed at 26°C. Five virgin female P0 were crossed to five male P0 per vial (for 1-2 vals) and flipped to new vials after one week. All emerged F1 were scored from both vials ∼15 days after initial pairing. F1 were scored for sex, genotype, and visible mutant phenotypes.

### Heat-shock induced mutant generation

To generate Cas9-mediated mutant *D. melanogaster*, the Cas9 strains were crossed with a gRNA strain to produce either double heterozygous F1 (*gRNA/+*; *Hsp70BbCas9/+*) or trans-heterozygous F1 (*Hsp70BbCas9/gRNA*) (both referred to as trans-heterozygous F1). Five virgin female P0 were crossed to five male P0 per vial and allowed to mate and lay eggs. Crosses at 18°C were allowed to seed a vial for four days, crosses at 21°C were allowed to seed a vial for three days, and crosses at 26°C were allowed to seed a vial for two days before P0 were flipped to new vials. All crosses performed at a specific temperature (18°C, 21°C, 26°C) utilized Cas9 stocks reared at the same temperature, unless otherwise specified. Vials with F1 embryos/first instar larvae were submerged in a 37°C water bath for heat shock durations of either 30 minutes, 1 hour, or 2 hours, then returned to their original rearing temperatures. All F1 were examined and scored for sex, genotype, and visible mutant phenotypes using a Leica M165FC fluorescent stereo microscope either after ∼27 days from initial pairing for 18°C rearing, ∼20 days for 21°C rearing, or ∼15 days for 26°C rearing.

### Sample collection and reverse transcription quantitative PCR (RT-qPCR)

All primers used for RT-qPCR are listed in the Reagents Table. Adult virgin female flies were used for RT-qPCR to maximize RNA extraction. Virgin females were collected and aged for 6-7 days. 7-10 females per condition were pooled together, heat shocked at 37°C as described above, and allowed to recover for 30 minutes at their original rearing temperature (i.e. 18°C, 21°C, 26°C). The flies were anesthetized, transferred to Eppendorf tubes on ice and immediately homogenized in QIAzol (Qiagen 79306) and stored at −20°C. RNA was extracted from samples following standard TRIzol/chloroform RNA extraction protocol (see Thermo Fisher Scientific TRIzol User Guide). RNA was quantified using a NanoDrop 2000 (Thermo Fisher Scientific) and diluted to 250 ng/μl, aliquoted, and stored at −80°C. 1μg of RNA was then treated with DNaseI (ThermoFisher Scientific #89836) to remove any DNA contamination, following a standard protocol. cDNA was synthesized using 5μL of DNaseI-treated RNA using RevertAid First Strand cDNA Synthesis Kit (ThermoFisher Scientific #K1622) and Oligo (dT)_18_ primers. qPCR was performed using 2x qPCRBIO SyGreen Blue Mix Separate-ROX (PCR Biosystems #PB20.17) on a LightCycler® 96 Instrument (Roche). cDNA samples from 21°C rearing with 2 hour heat shock samples were serially diluted to generate standard curves for each amplified gene fragment and to test primer performance (Figure S2A-B). Undiluted samples (d0) were used for relative quantification to account for possible low Cas9 expression levels for some samples. RT-qPCR reactions (20μL) were ran following 2x qPCRBIO SyGreen Blue Mix Separate-ROX (PCR Biosystems #PB20.17) protocol with 2μL of sample. Negative controls were run with 2μL of nuclease-free water. At least three technical replicates were run per plate for each of three biological replicates. Data from 2-3 technical replicates for all three biological replicates were analyzed in LightCycler® 96 software (Roche), Google Sheets, and GraphPad Prism 9. Relative ratios of *Cas9* RNA levels normalized to *RpL32* were calculated using E_*RpL32*_ ^RpL32_Ct^/E_*Cas9*_ ^Cas9_Ct^. Normalized expression ratio (or fold change) was also calculated using the ΔΔCt method.

### Genotyping target loci

Control flies (*w*^*1118*^ and heterozygous flies) and trans-heterozygous flies with or without visible mutant phenotypes (or lethal pupae) were separately collected in Eppendorf tubes and frozen at −20°C. Genomic DNA was extracted from individual flies following a standard DNA squish protocol. In short, single flies were homogenized in 30μL Tris-EDTA buffer pH 8.0 with 5M NaCl and Proteinase-K (proK) Qiagen solution (from Qiagen #69504). Samples were incubated at 37°C for 30 minutes and then at 95°C for four minutes. Samples were either frozen immediately and/or used for PCR amplification. PCR amplification of all target regions was performed using LongAmp Taq 2X Mastermix (NEB #M0287). Amplicon size was verified using gel electrophoresis, and amplicons were purified using QIAquick PCR Purification Kit (Qiagen # 28104) before Sanger sequencing through Retrogen Inc. Primers used can be found in the Reagents Table. All sequence file chromatograms were verified and aligned to the following NCBI Reference Sequences: NC_004353.4 (*ey*), NT_033777.3 (*Ser*) (Larkin *et al*. 2021) using SnapGene® 4.

### Microscopy

Representative images of mutant phenotypes and lethal pupae were obtained using light microscopy on a Leica M165FC fluorescent stereomicroscope equipped with the Leica DMC2900 camera. Scale bars were designed manually with a ruler in each original image. Images were then processed for ease of viewing and assembled with Adobe Photoshop.

### Statistical analysis

Percentage of mutant formation for heat-shock induced mutant generation (Figures 3, 4, 5) was calculated using the total number of flies scored for a genotype/condition over the total number of flies scored (combining multiple vials) to minimize any effect by variability across vials or crosses. This percentage was reported along with the standard error of the mean. Significance was calculated using unpaired two-tailed t-test with Welch’s correction using GraphPad Prism9. Significance for all RT-qPCR data was calculated using unpaired two-tailed t-test with Welch’s correction using GraphPad Prism9.

## Results

### Construct design and *Drosophila* transgenesis

The *Cas9* (*Csn1*) endonuclease from the *Streptococcus pyogenes* Type II CRISPR/Cas system was utilized downstream the *Drosophila melanogaster Hsp70Bb* promoter followed by a T2A sequence and enhanced GFP (eGFP) to provide heat-shock inducible expression of *Cas9* (Kandul *et al*. 2021). *S. pyogenes Cas9* has been widely used to induce genome modification in *Drosophila melanogaster* (Bassett et al. 2013; Gratz et al. 2013, 2014; Yu et al. 2013; Xue et al. 2014) as well as many other model organisms (Wang *et al*. 2016). The *Drosophila Hsp70Bb* promoter has been widely used for inducible gene expression and has been reported to have leaky expression, where the downstream elements are expressed without incubation at the inducible heat-shock temperature (Curtin *et al*. 2008; Naidoo and Young 2012; Akmammedov *et al*. 2017). Suppression of leaky expression of *lacZ* under control of the *Hsp70Bb* promoter was achieved through the addition of upstream *bithoraxoid* (*bxd*) Polycomb response elements (PREs) with insulating gypsy elements (Simon *et al*. 1993; Akmammedov *et al*. 2017). Given this strong suppression of promoter activity, we sought to utilize PREs to minimize leakiness of *Cas9* under *Hsp70Bb* promoter control. The PRE repeats and gypsy insulator elements were cloned into the original Hsp70Bb-Cas9-T2A-eGFP vector and used for site-specific ϕC31 integration to generate *PRE-Hsp70BbCas9* transgenic *D. melanogaster* (see Materials and Methods and Figure S1) for characterization and comparison against *Hsp70BbCas9* transgenic *D. melanogaster*.

### Polycomb response elements (PREs) decrease relative *Cas9* transcript levels

Reverse transcription quantitative PCR (RT-qPCR) was used to assess the relative levels of *Cas9* transcripts with and without upstream PREs at multiple heat shock durations and *D. melanogaster* rearing temperatures (Figure 1A). Levels of *Cas9* transcripts were assessed relative to the housekeeping gene *Ribosomal protein L32* (*RpL32*) (Figure S2A-B). *ATP synthase, coupling factor 6* (*ATPsynCF6*) was not used as a reference as done previously (Akmammedov *et al*. 2017; Kandul *et al*. 2021) since there was no significant difference in average Cq values between samples from *PRE-Hsp70BbCas9* 18°C with no heat shock (hypothesized low promoter activity) and samples from *Hsp70BbCas9* at 21°C rearing after a 2 hour heat shock (hypothesized high promoter activity) (Figure S2C).

**Figure 1.**
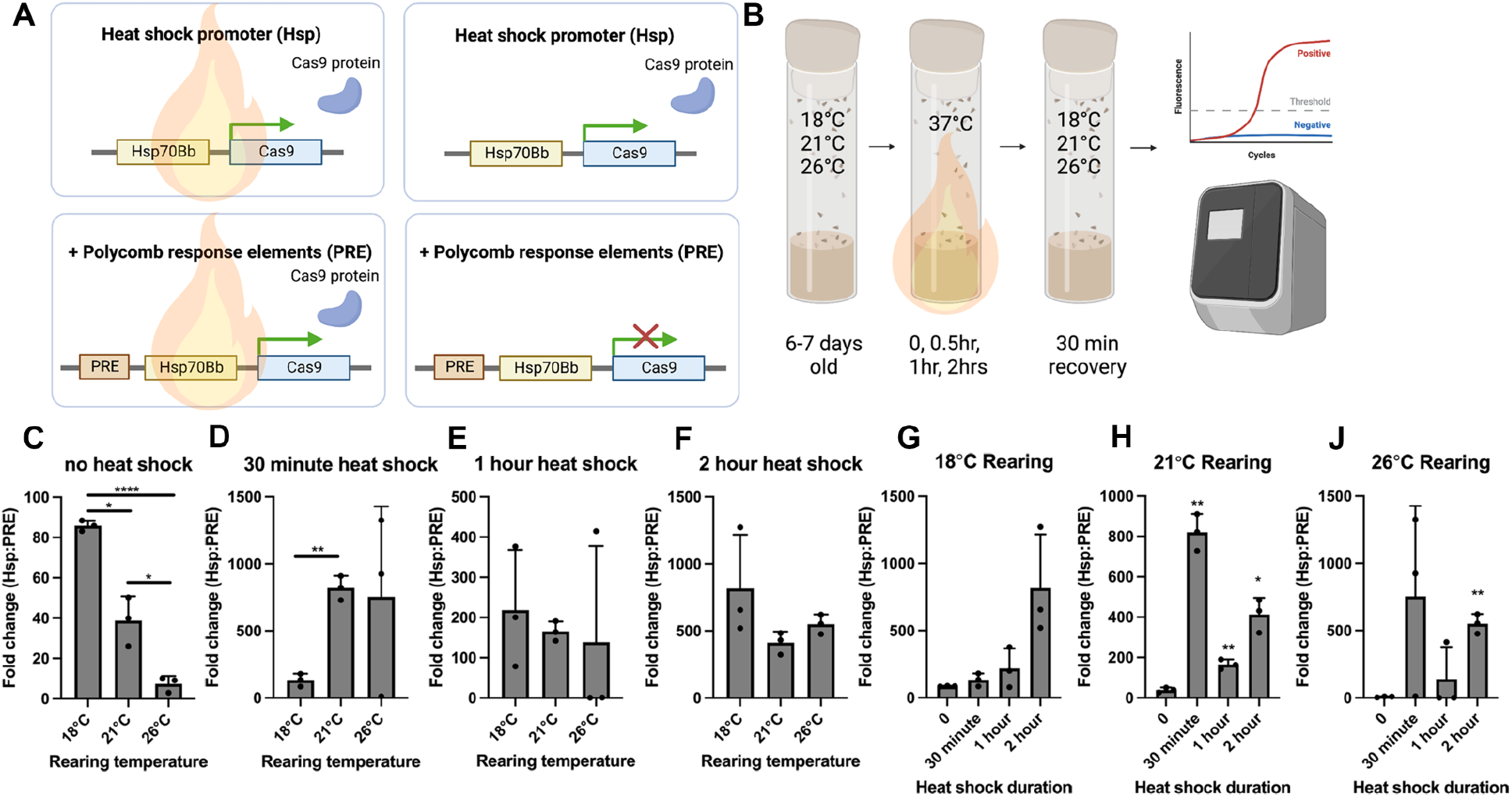
Fold change of *Cas9* RNA levels by RT-qPCR comparing the lack of upstream Polycomb response elements (PRE) to having PREs, considering varying heat shock durations and rearing temperatures. A) Conceptual use of Polycomb response elements (PREs) to suppress leaky expression of *Cas9* under heat shock-inducible *Drosophila melanogaster Hsp70Bb* promoter. Created with BioRender.com. B) Methodology for sample collection and RT-qPCR. Created with BioRender.com. Normalized expression ratio (fold change, calculated using ΔΔCt method) of *Cas9* transcripts for *Hsp70BbCas9*: *PRE-Hsp70BbCas9* (Hsp:PRE) plotted against C-F) rearing temperature (per heat shock duration) and against G-J) heat shock duration (per rearing temperature). C-F and G-J are the same data, replotted for visualization purposes. Error bars = standard deviation. Significance was calculated using unpaired two-tailed t-test with Welch’s correction. ns (or unlabeled) = P > 0.05, * = P ≤ 0.05, ** = P ≤ 0.01, **** = P ≤ 0.0001. Statistics for G-J were calculated for heat shock duration (30 minutes, 1 hour, 2 hours) compared to no heat shock.

Relative levels of *Cas9* transcripts were compared for *Hsp70BbCas9* and *PRE-Hsp70BbCas9* flies at different heat shock durations and rearing temperatures (Figure S2D-K). It is important to note that *Cas9* transcripts are present after no heat shock from *PRE-Hsp70BbCas9* flies (Figure S2D). After no heat shock, significant differences in relative *Cas9* transcript levels were found at 18°C and 21°C rearing temperatures, but not at 26°C, suggesting that higher rearing temperatures may negate any effect of the PREs on *Hsp70Bb* promoter activity (Figure S2D). This suggests that PREs do not completely suppress *Hsp70BbCas9* leaky expression in this system. Average relative levels of *Cas9* transcripts generally increased with increasing heat shock duration at all rearing temperatures for *Hsp70BbCas9* flies, with significant increases compared to no heat shock for 18°C rearing after a 30 minute heat shock (0.74 ± 0.26, p = 0.0388; vs no heat shock = 0.0035 ± 0.00010) (Figure S2E), 21°C rearing for all heat shock durations (30 minute = 0.77 ± 0.083, p = 0.0039; 1 hour = 0.87 ± 0.12, p = 0.0066; 2 hours = 1.26 ± 0.24, p = 0.0121; vs no heat shock = 0.00081 ± 0.00025) (Figure S2F), and 26°C rearing after a 2 hour heat shock (0.57 ± 0.074, p = 0.0056; vs no heat shock = 0.00023 ± 0.00012) (Figure S2G). High variability was observed after heat shock for 18°C and 26°C reared *Hsp70BbCas9* flies. Relative levels of *Cas9* transcripts increased significantly in *PRE-Hsp70BbCas9* flies after all heat shock durations at 18°C rearing (30 minute = 0.0063 ± 0.0013, p = 0.0137; 1 hour = 0.0058 ± 0.00069, p = 0.0048; 2 hours = 0.0034 ± 0.00038, p = 0.0043; vs no heat shock = 0.000047 ± 0.000017) (Figure S2H), and after both 1 hour and 2 hour heat shocks at 21°C rearing (1 hour = 0.0059 ± 0.00048, p = 0.0022; 2 hours = 0.0036 ± 0.00082, p = 0.0176; vs no heat shock = 0.000023 ± 0.0000015) (Figure S2J) as well as at 26°C rearing (1 hour = 0.0019 ± 0.00055, p = 0.0286; 2 hours = 0.0012 ± 0.00033, p = 0.0237; vs no heat shock = 0.000034 ± 0.000014) (Figure S2K). Interestingly, average relative levels of *Cas9* transcripts from *PRE-Hsp70BbCas9* samples after a 2 hour heat shock were lower compared to after a 1 hour heat shock (Figure S2H-K). This may be indicative of the rapid suppression of *Cas9* expression during the 30 minute recovery period after heat shock (Akmammedov *et al*. 2017).

Fold change in *Cas9* expression was calculated for these RT-qPCR data using the ΔΔCt method to assess the fold difference in *Cas9* expression in *Hsp70BbCas9* normalized to *PRE-Hsp70BbCas9* flies. Without heat shock, *Hsp70BbCas9* animals had a 85.85 ± 2.63 fold higher expression of *Cas9* at 18°C rearing temperature than *PRE-Hsp70BbCas9* animals (Figure 1C). As rearing temperature increased, this fold change in expression decreased (38.80 ± 12.11 for 21°C and 7.41 ± 3.90 for 26°C) (Figure 1C). After heat shock, the average fold change in expression exceeded 130 (133.01 ± 47.77 for 30 minute heat shock, 18°C rearing, see Figure 1C) with the highest average fold change at 820.49 for 30 minute heat shock, 21°C rearing (Figure 1D). After 1 hour or 2 hour heat shocks, no significant difference in fold change in expression of *Cas9* was observed across all rearing temperatures (Figure 1E-F). Revisualizing these data as fold change by heat shock duration per rearing temperature reveals large differences in average fold change after 30 minute heat shock compared to no heat shock at 21°C and 26°C rearing temperatures (Figure 1H-J). The largest average fold change difference for 18°C rearing was after a 2 hour heat shock (Figure 1G). Significant differences in fold change compared to no heat shock conditions were found for 21°C rearing after all heat shock durations (30 minutes = 820.49 ± 90.54, p = 0.0039; 1 hour = 165.54 ± 24.47, p = 0.0044; 2 hour = 412.57 ± 82.94, p = 0.0145; vs no heat shock = 38.8 ± 12.11) (Figure 1H), and for 26°C rearing after a 2 hour heat shock (550.50 ± 72.58, p = 0.0058; vs no heat shock = 7.41 ± 3.90) (Figure 1J). Overall, rearing *PRE-Hsp70BbCas9* flies at either 18°C or 21°C and using either a 2 hour or 30 minute heat shock, respectively, could allow for both minimization of leaky expression of *Cas9* without heat shock and maximization of *Cas9* expression after heat shock.

### Selection of gene targets and confirmation of mutant phenotypes

Given the significant decrease in, but not complete suppression of, *Cas9* transcripts with Polycomb response elements (PREs) upstream the *Hsp70Bb* promoter, we sought to characterize to what extent these relatively low levels of Cas9 are able to induce mosaic mutant phenotypes. To further characterize the ability of the the inducible promoter to generate functional levels of Cas9, *PRE-Hsp70BbCas9 D. melanogaster* transgenic line was compared to *Hsp70BbCas9* for their ability to generate F1 mutant phenotypes at different rearing temperatures (18°C, 21°C, and 26°C) and varying heat shock durations (no heat shock, 30 minutes, 1 hour, or 2 hours) (Figure 2A). Three genes with established guide RNA (gRNA) *D. melanogaster* stocks (*sgRNA:w, dgRNA:ey, dgRNA:Ser*) were used to characterize mutant phenotype generation (Figure 2A, Reagents Table). *sgRNA:w* was used since targeting the *white* gene often results in a prominent, visible phenotype, allowing for ease of identification while also allowing for a description of a high-efficacy gRNA. *sgRNA:ey*, targeting the *eyeless* gene also results in a prominent, visible phenotype, but was chosen given its location on the highly heterochromatic 4th, or dot, chromosome, as an example for a low-efficacy gRNA stock. *sgRNA:Ser* was chosen to target *Serrate*, a dominant gene with a visible and often lethal phenotype.

**Figure 2.**
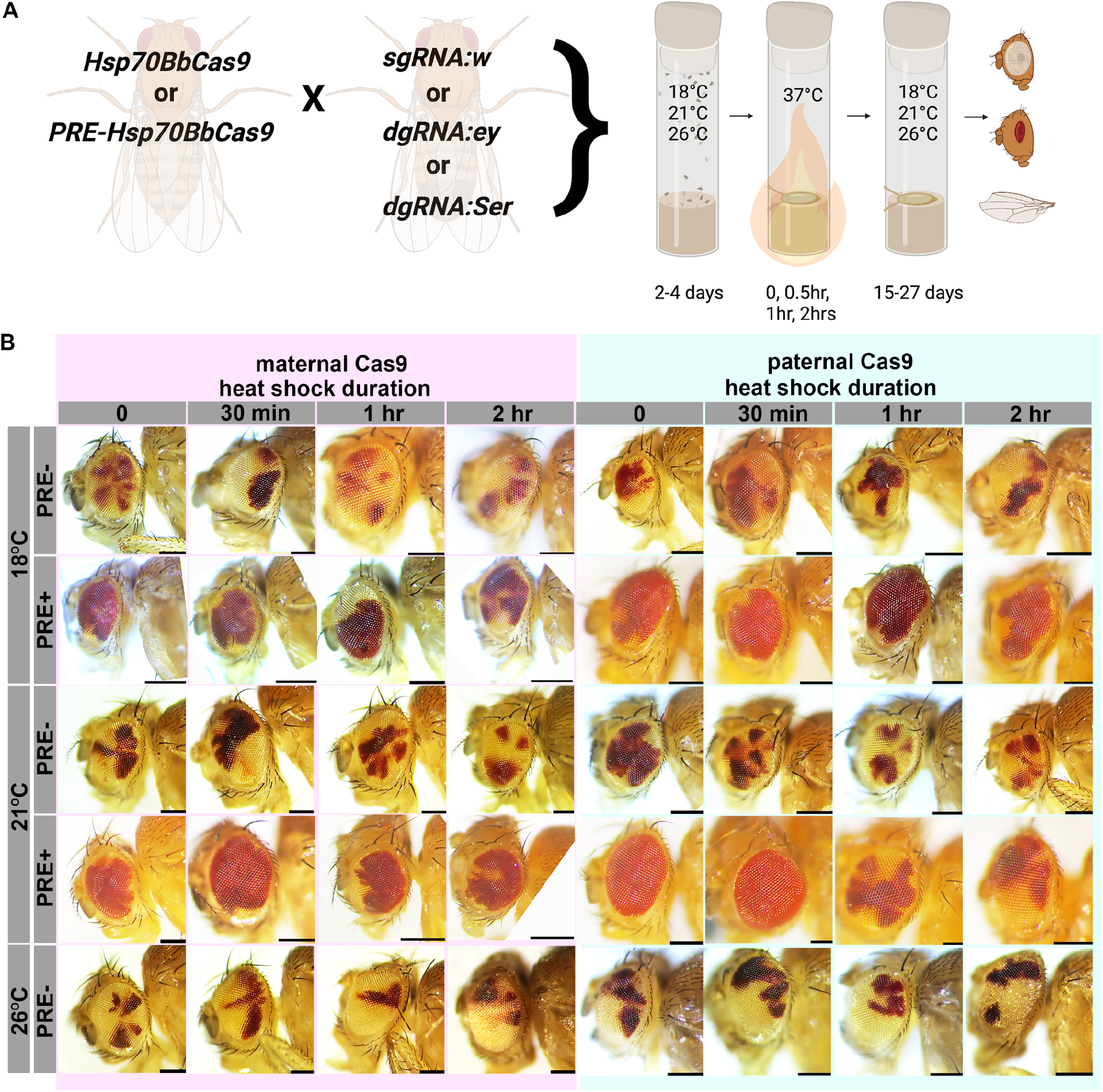
Polycomb response elements (PREs) decrease the severity of the *white* mutant phenotype. A) General schematic of genetic crosses and heat shock to generate *Cas9; gRNA* trans-heterozygotes. Created with BioRender.com. B) Representative images of *white* eye mutant phenotype variegation with and without PREs at multiple rearing temperatures (18°C, 21°C, or 26°C) and heat shock durations (0 or no heat shock, 30 minute heat shock, 1 hour heat shock, or 2 hour heat shock). The left four columns (highlighted in magenta) are F1 from maternal *Hsp70BbCas9* or *PRE-Hsp70BbCas9* and the right four columns (highlighted in turquoise) are F1 from paternal *Hsp70BbCas9* or *PRE-Hsp70BbCas9*. PRE+ flies reared at 26°C were not scored or imaged due to suppressed *mini-white*. All scale bars = 250μm.

Mutant phenotypes were generated with constitutively expressed *Cas9* stocks, nanos-Cas9 (*nosCas9*) and vasa-Cas9 (*vasaCas9*), as controls. When crossed with *sgRNA:w, nosCas9* generated a low frequency of variegated eyes in the F1 (maternal Cas9: 5.71% of F1 females and 6.38% F1 males; paternal Cas9: 0% for both F1 females and males) compared to *vasaCas9* (maternal Cas9: 100.0 ± 0.0% of F1 females, males were not scored; paternal Cas9: 100.0% for both F1 females and males) (Figure S3A). *nosCas9* also generated milder phenotypes (FigureS3B-C) compared to *vasaCas9* (Figure S3D-E). When crossed with *dgRNA:ey, nosCas9* generated zero mutants (maternal Cas9: 0% of both F1 females and males; paternal Cas9: 0% of both F1 females and males) (Figure S3G) compared to *vasaCas9* (maternal Cas9: 16.67 ± 23.57% of F1 females and 33.33 ± 23.57% of F1 males; paternal Cas9: 30.0 ± 42.43% of F1 females and 28.57 ± 40.41% F1 males) (Figure S3F). Maternal *vasaCas9* produced slightly more mutant progeny than paternal *vasaCas9*, however, the severity of the mutant phenotype was often similar (Figure S3H,K) with rare full phenotypes (Figure S3J). When crossed with *dgRNA:Ser, nosCas9* was able to generate viable F1 whereas *vasaCas9* generated lethal phenotypes (Figure S3L). *nosCas9* was further able to generate *Serrate* mutant phenotypes when inherited maternally (3.33% of F1 females and 4.76% of F1 males), but not paternally (0% of F1 females and F1 males) (Figure S3L). These results provide a range of mutant rate formation, where *nosCas9* generally generated lower rates of or less severe phenotypes compared to *vasaCas9*. Simultaneously, these data allow for comparison to the rate of mutant phenotype formation generated by inducible *Cas9* expression.

**Figure 3.**
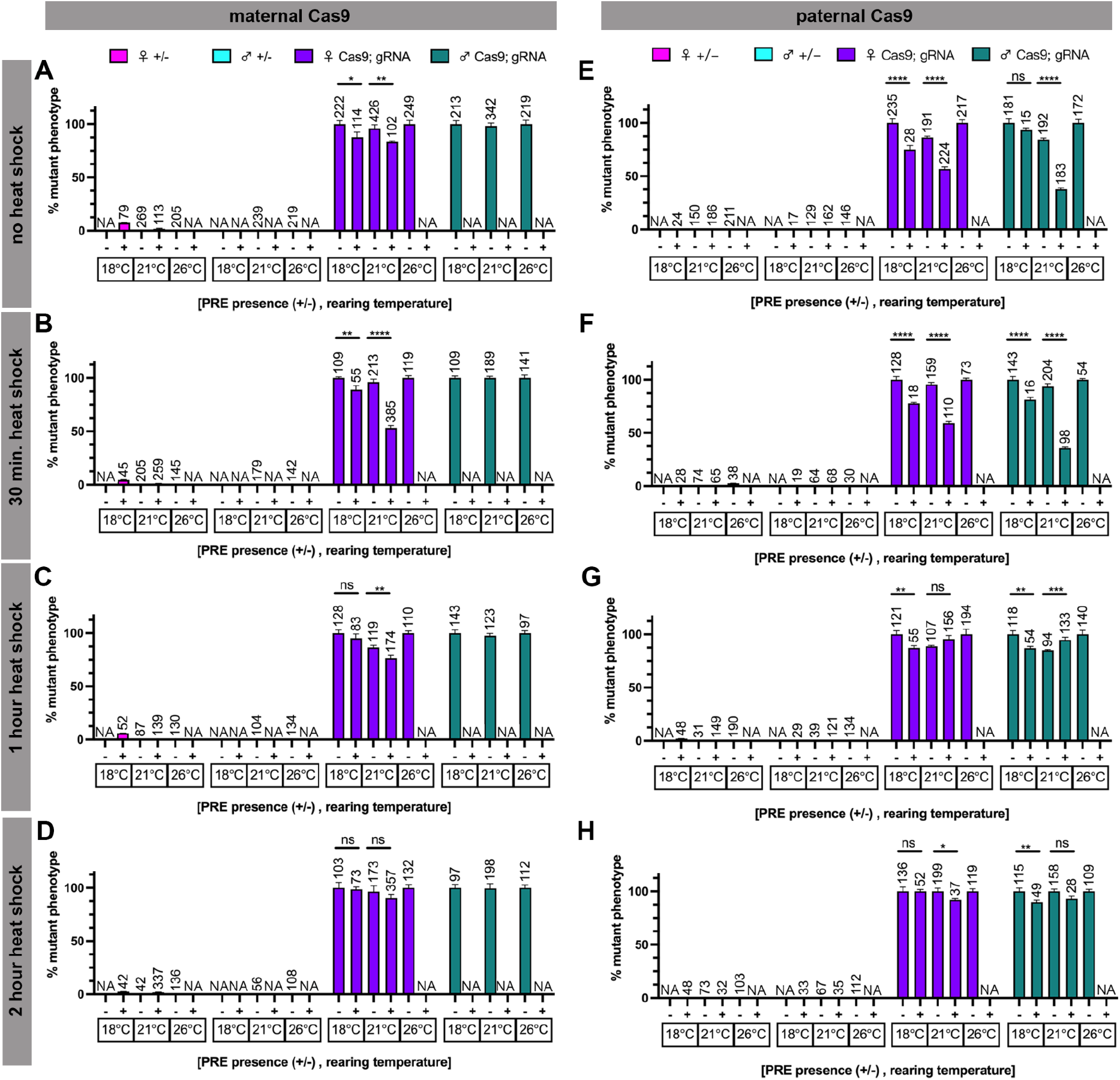
Polycomb response elements (PRE) decrease rates of Cas9-induced *white* eye phenotypes. Rate of mutant phenotype generation with heat shock inducible *Cas9* (*Hsp70BbCas9*) and *sgRNA:w* with and without upstream Polycomb response elements (+/-PRE, *Hsp70BbCas9* compared to *PRE-Hsp70BbCas9*), considering heat shock duration, parental contribution, and rearing temperature. Percentage of mutant phenotype in F1 with A-D) maternal contribution of Cas9 after A) no heat shock, B) 30 minute heat shock, C) 1 hour heat shock, D) 2 hour heat shock. Percentage of mutant phenotype in F1 with E-H) paternal contribution of Cas9 after E) no heat shock, F) 30 minute heat shock, G) 1 hour heat shock, and H) 2 hour heat shock. Magenta = heterozygous female controls, turquoise = heterozygous male controls, purple = trans-heterozygous females, green = trans-heterozygous males. n = number of flies scored (listed above corresponding bar). Error bars = standard error of the mean. NA = not scored, see Results for details. Y-axis extends above 100% to account for error bars. Significance was not calculated for heterozygous controls. Significance was calculated using unpaired two-tailed t-test with Welch’s correction. ns = P > 0.05, * = P ≤ 0.05, ** = P ≤ 0.01, *** = P ≤ 0.001, **** = P ≤ 0.0001.

### Polycomb response elements (PREs) reduce the severity of *white* mutant phenotype induced by *Hsp70BbCas9*

Trans-heterozygous *sgRNA:w; Cas9* F1 flies with or without PREs were scored for the presence of any *white* mutant phenotype after heat shock. The rate of *white* mutant phenotype formation in *sgRNA:w; Cas9* trans-heterozygous flies was high across all conditions (Figure 2-3) (all but two conditions were over 50%), and closer to rates of mutant phenotype formation by *vasaCas9* controls (maternal Cas9: 100 ± 0% F1 females; paternal Cas9: 100% F1 females and 100% F1 males) compared to *nosCas9* controls (maternal Cas9: 5.71% F1 females and 6.38% F1 males; paternal Cas9: 0% F1 females and 0% F1 males) (Figure S3A). All heat shock durations at both 18°C and 26°C rearing temperatures for maternal and paternal *Hsp70BbCas9* (PRE-trans-heterozygotes) averaged 100% visible mutant phenotypes (Figure 3). The rate of visible mutant phenotypes varied for 21°C rearing temperatures for *Hsp70BbCas9* F1 trans-heterozygous flies, but did not drop below 84% (Figure 3). The reason for this discrepancy between rearing temperatures remains unknown. These results suggest that there is no obvious or consistent difference between *Hsp70BbCas9* maternal and paternal inheritance in combination with *sgRNA:w*.

Presence of PREs upstream of the *Hsp70Bb* promoter significantly decreased rates of *white* mutant phenotypes for most conditions tested (parental Cas9 inheritance, rearing temperature, and heat shock duration) (Figure 3). For both maternal and paternal Cas9 inheritance, as the heat shock duration increased, differences between mutant phenotype formation in female F1 lost significance. After a 2 hour heat shock, there was no significant difference in percent mutant phenotype of female F1 with maternal Cas9 with or without PREs at 18°C rearing (PRE+: 98.63 ± 2.19% vs PRE-: 100 ± 5.08%) or 21°C rearing (PRE+: 90.20 ± 3.80% vs PRE-: 96.53 ± 5.39%), nor a significant difference in percent mutant phenotype of F1 with paternal Cas9 with or without PREs at 18°C rearing for female F1 (PRE+: 100 ± 1.80% vs PRE-: 100 ± 3.97%) or at 21°C rearing for male F1 (PRE+: 92.86 ± 2.90% vs PRE-: 100 ± 2.10%) (Figure 3). This suggests that the longer the heat shock duration, the less effective PREs become in suppressing functional levels of *Cas9* transcripts in the presence of a highly efficacious gRNA.

Representative images of trans-heterozygous flies were collected to qualitatively evaluate severity of *white* variegation since the binary calls of our quantification of the rate of mutant formation does not represent the extent of variegation. The majority of the *white* mutant trans-heterozygous flies without PREs (*sgRNA:w; Hsp70BbCas9*) (Figure 2, rows 1, 3, and 5) show strong variegation phenotypes, regardless of parental Cas9 contribution (maternal vs paternal), rearing temperature, or heat shock duration. Trans-heterozygous flies with PREs (*sgRNA:w; PRE-Hsp70BbCas9*) show a range of variegation phenotypes: some flies show no variegation (e.g. paternal Cas9, 21°C rearing, no heat shock and 30 minutes heat shock), and the severity of variegation appears to increase with longer heat shocks at all rearing temperatures (Figure 2, rows 2 and 4). *sgRNA:w; PRE-Hsp70BbCas9* trans-heterozygous flies reared at 26°C were not scored due to PRE suppression of the *mini-white* transgenic marker (Kassis *et al*. 1991; Akmammedov *et al*. 2017) to avoid any uncertainty regarding the presence of the PRE-Hsp70Bb-Cas9 transgene (labeled as NA in Figure 3). Heterozygous controls at 18°C were often not included (labeled as NA in Figure 3); the *CyO* balancer curly wing phenotype often appeared suppressed in *CyO; Hsp70BbCas9* heterozygous F1 (not scored). This suppressed curly wing phenotype is most likely due to reduced expression of *Curly* (*Cy*) at temperatures below 19°C (Pavelka *et al*. 1996). Given the difficulty to distinguish between heterozygous and trans-heterozygous F1 flies, only homozygous *sgRNA:w* P0 were used to generate trans-heterozygous *sgRNA:w; Hsp70BbCas9* flies at 18°C. Finally, *sgRNA:w; PRE-Hsp70BbCas9* trans-heterozygous males with maternally inherited Cas9 were not scored due to difficulty scoring the low contrasted (and PRE-suppressed) *mini-white* transgene variegation in a *white* mutant background (Kassis *et al*. 1991; Akmammedov *et al*. 2017). Overall, PREs upstream *Hsp70BbCas9* are able to decrease the severity of *white* mutant variegation for most conditions tested.

Individual trans-heterozygous *sgRNA:w; Hsp70BbCas9* flies were genotyped and compared to heterozygous controls to identify the presence of targeted DNA mutations by Cas9. Since F1 from *sgRNA:w* crosses were heterogeneous for *white* alleles which can not be separated by Sanger sequencing, the *mini-white* transgene was sequenced for genotyping purposes. Cas9 was able to induce DNA mutations in trans-heterozygous *sgRNA:w; Hsp70BbCas9* flies with mutant phenotypes when inherited maternally (Figure S4C) or paternally (Figure S4E), suggesting that scoring mutant phenotype rate can act as a proxy for Cas9 function.

### PREs upstream *Hsp70BbCas9* reduce the efficacy of already low-efficiency guide RNAs targeting *eyeless*

The rate of *eyeless* (*ey*) mutant phenotype formation was very low across all conditions (Figure 4F-N) (under 10%), consistent with low mutant phenotype formation with constitutive *nosCas9* and *vasaCas9* controls (Figure S3G). Rearing temperature did not change the low rate of *ey* mutant phenotype formation with no heat shock (Figure 4F, K). All mutant phenotype rates were relatively low with the highest rate of mutant phenotype formation seen at 26°C rearing temperature for F1 females inheriting maternal *Hsp70BbCas9* (4.62 ± 0.18%) and F1 males inheriting paternal *Hsp70BbCas9* (8.22 ± 0.23%) both after a 2 hour heat shock (Figure 4F-N). The majority of heat-shock induced *ey* phenotypes observed were mild, similar to Figure 4A, Figure 4C, and Figure 4E; on rare occasions a large portion or the majority of the eye would not develop, as in Figure 4B and Figure 4D (severity of mutant phenotype not quantified). Regardless of low mutant phenotype generation, presence of PREs significantly decreased the rate of mutant phenotype in F1 in every condition where mutants were identified (Figure 4F-N). In fact, *ey* mutants were only identified in *dgRNA:ey; PRE-Hsp70BbCas9* trans-heterozygous F1 under one condition (female F1 at 18°C rearing and paternal Cas9: 1.39 ± 0.13%) (Figure 4K).

**Figure 4.**
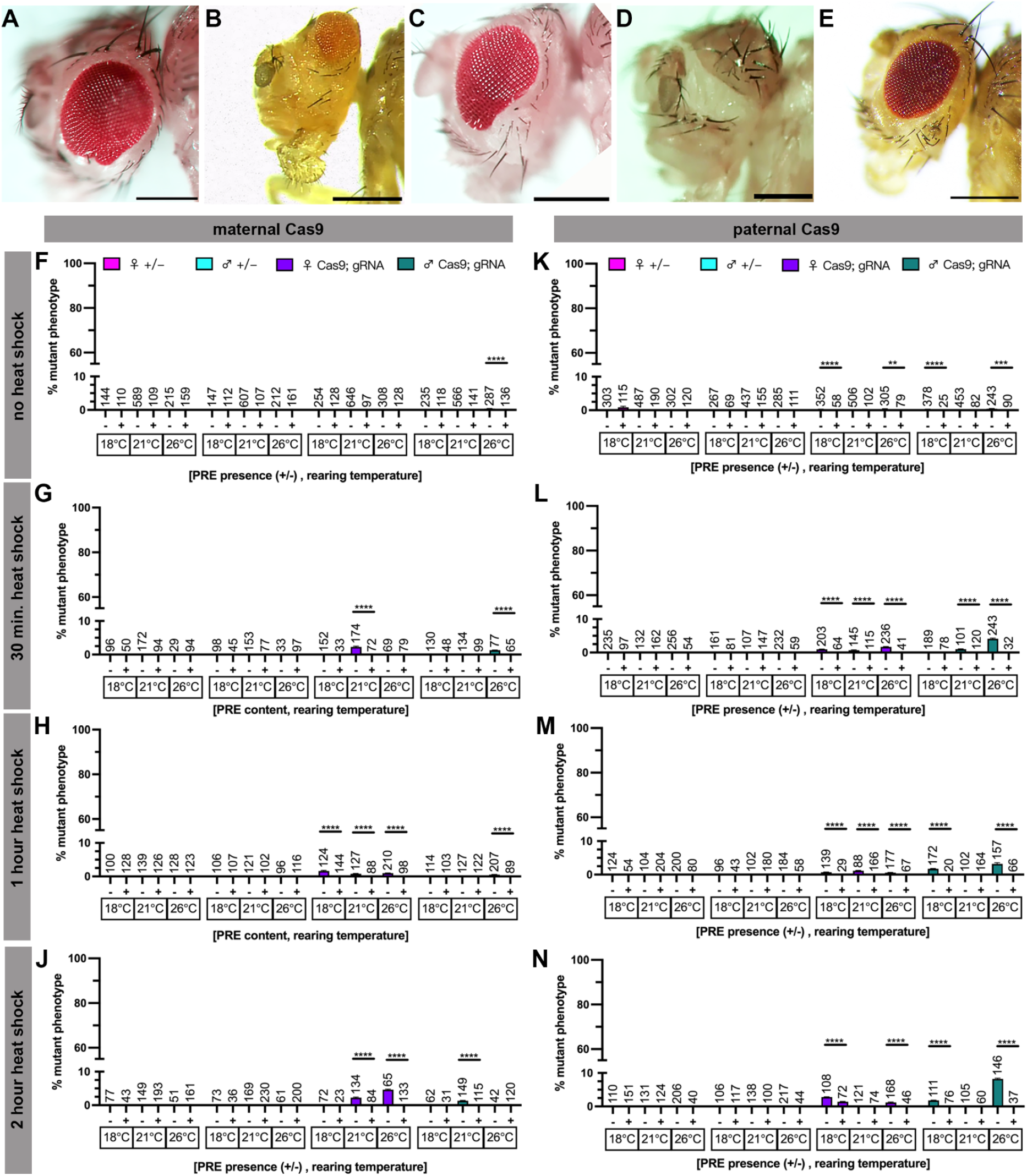
PREs further reduce a rare eyeless phenotype. A-E) Representative images of the phenotypic range of induced *eyeless* phenotypes with A-B) maternal *Hsp70BbCas9*, C-D) paternal *Hsp70BbCas9*, and E) maternal *PRE-Hsp70BbCas9*. All scale bars = 250μm. F-N) Rate of mutant phenotype generation with heat shock inducible *Cas9* (*Hsp70BbCas9*) and *dgRNA:ey* with and without upstream Polycomb response elements (+/-PRE), considering heat shock duration, Cas9 contribution, and rearing temperature. Percentage of mutant phenotype in F1 with F-J) maternal contribution of Cas9 after F) no heat shock, G) 30 minute heat shock, H) 1 hour heat shock, J) 2 hour heat shock. Percentage of mutant phenotype in F1 with K-N) paternal contribution of Cas9 after K) no heat shock, L) 30 minute heat shock, M) 1 hour heat shock, and N) 2 hour heat shock. Magenta = heterozygous female controls, turquoise = heterozygous male controls, purple = trans-heterozygous females, green = trans-heterozygous males. n = number of flies scored (listed above corresponding bar). Error bars = standard error of the mean. Significance was not calculated for heterozygous controls. Unlabeled = 0 ± 0% for both values (and no significance calculated). Significance was calculated using unpaired two-tailed t-test with Welch’s correction. ** = P ≤ 0.01, *** = P ≤ 0.001, **** = P ≤ 0.0001.

Individual flies were genotyped using Sanger sequencing to identify the presence of targeted DNA mutations by Cas9 in flies with or without a mutant phenotype. All individuals sequenced showed no DNA modifications compared to controls for *ey* gRNA target 1 (Figure S5, left column). Regardless of phenotype status, trans-heterozygous individuals sequenced showed low levels of secondary peaks adjacent to the *ey* gRNA target 2 site (Figure S5D, F, H, K, right column) compared to most heterozygous controls (Figure S5C, E, G, J). Paternal *Hsp70BbCas9* heterozygous control however revealed similar low levels of secondary peaks (Figure S5E). The reason for this subtle sequence difference remains unclear, however, given the low rates of mutant phenotype generation and low severity of mutant phenotypes suggests that the majority of DNA sequences present may have been wild type, obscuring any mutant genotype. We suspect that PCR amplicons of the wild type sequences were in high abundance compared to mutant sequences given that sequencing of individual F1 from *vasaCas9* crosses revealed multiple peaks at both *ey* gRNA target 1 and gRNA target 2 sites (Figure S6E and G).

### PREs upstream *Hsp70BbCas9* allow for the generation of viable *Serrate* mutants

Mutations of *Serrate* by *Hsp70BbCas9* resulted in complete lethality (100%) of trans-heterozygous flies regardless of heat shock duration or rearing temperature (Figure 5F-N). All transheterozygous flies appeared to die during pupal development (Figure 5A and B, unscored). Individual lethal pupae were genotyped using Sanger sequencing and found to have sequence mutations at both *Ser* gRNA target 1 and gRNA target 2 sites compared to adult heterozygous controls (Figure S7). Addition of PREs allowed for viable trans-heterozygotes and allowed for formation of visible mutant phenotypes (Figure 5D and E) for most conditions (Figure 5F-N).

**Figure 5.**
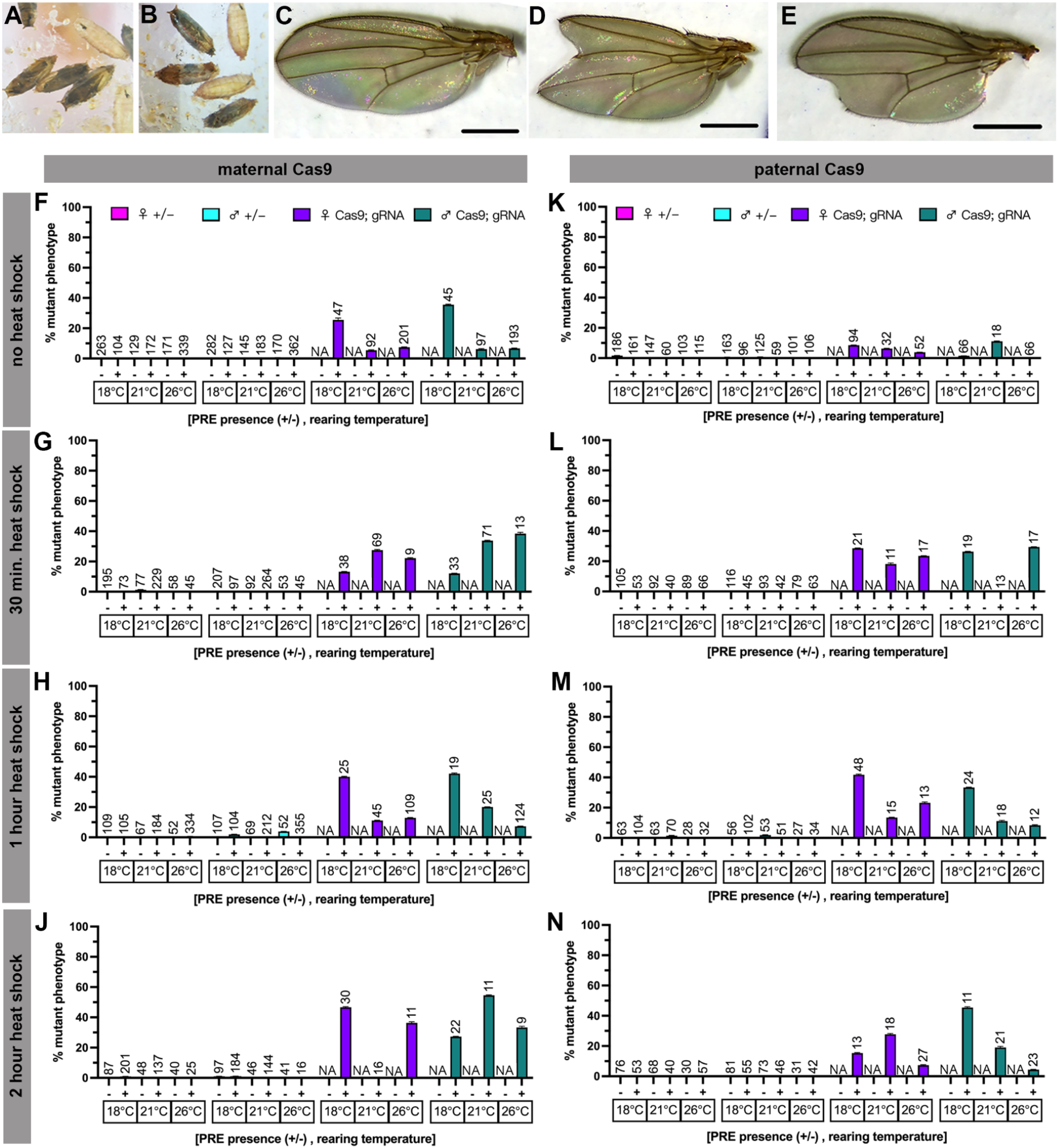
PRE elements allow for viable *Serrate* phenotypes in an otherwise lethal genotype. Representative images of induced *Serrate* mutant phenotypes with *dgRNA:Ser*. A-B) Lethal pupae from A) maternal *Hsp70BbCas9*, and B) paternal *Hsp70BbCas9*. C-E) *PRE-Hsp70BbCas9*/*dgRNA:Ser* trans-heterozygotes wing examples with maternal *PRE-Hsp70BbCas9* at 26°C rearing with no heat shock C) with no serration, and D,E) with serration. All scale bars = 500μm. Percentage of mutant phenotype generation with heat shock inducible *Cas9* (*Hsp70BbCas9*) and *dgRNA:Ser* with and without upstream polycomb response elements (+/-PRE), considering heat shock duration, Cas9 contribution, and rearing temperature. F-J) Maternal contribution of Cas9 after F) no heat shock, G) 30 minute heat shock, H) 1 hour heat shock, J) 2 hour heat shock. K-N) Paternal contribution of Cas9 after K) no heat shock, L) 30 minute heat shock, M) 1 hour heat shock, and N) 2 hour heat shock. Magenta = heterozygous female controls, turquoise = heterozygous male controls, purple = trans-heterozygous females, green = trans-heterozygous males. n = number of flies scored (listed above corresponding bar). Error bars = standard error of the mean. Significance was not calculated for heterozygous controls. Significance was not calculated for trans-heterozygous flies since all *Hsp70BbCas9*/*dgRNA:Ser* (PRE -) flies died during metamorphosis.

To verify DNA mutations in viable trans-heterozygotes, individual *dgRNA:Ser; PRE-Hsp70BbCas9* flies with or without a visible phenotype were genotyped by Sanger sequencing (Figure S8). Individual F1 with maternal *PRE-Hsp70BbCas9* without serrated wings, revealed nucleotide changes only at gRNA target 1 (Figure S8D), suggesting either differences in the severity of different mutant alleles, or insufficient mosaicism to induce a visible phenotype. F1 with maternal *PRE-Hsp70BbCas9* with serrated wings revealed nucleotide changes at both gRNA targets (Figure S8E). Individual F1 with paternal *PRE-Hsp70BbCas9* showed no nucleotide changes at both gRNA target sites regardless of the presence of a visible serrated wing (Figure S8G and H). This either suggests that paternal *PRE-Hsp70BbCas9* resulted in fewer mutant cells in mosaic animals compared to maternal Cas9, or false positives (general wing damage) were scored as *Serrate* mutants (Figure 5, see heterozygous control values).

Pupal lethality of *dgRNA:Ser; vasaCas9* trans-heterozygotes corresponded to drastic sequence differences at gRNA target 1 and 2 for both maternal (Figure S9G and L, respectively) and paternal *vasaCas9* (Figure S9H and M, respectively). Sanger sequencing of *dgRNA:Ser; nosCas9* trans-heterozygotes revealed drastic nucleotide changes between both gRNA target sites with maternal *nosCas9* (Figure S9C), seemingly corresponding to the reported visible mutant phenotype (3.33% of F1 females and 4.76% of F1 males) (Figure S3L). No nucleotide differences were observed for paternal *nosCas9* (FigureS9D) corresponding to a rate of 0% mutant phenotype formation (Figure S3L).

## Discussion

### Polycomb response elements (PREs) significantly reduce leakiness of *Cas9* without heat shock

Here we characterize the ability of Polycomb response elements (PREs) to suppress heat-shock induced *Cas9* expression and indirectly, the ability to limit Cas9 functionality. Our results corroborate previous reports characterizing the ability of PREs to significantly suppress leaky expression of the *Drosophila Hsp70Bb* promoter (Akmammedov *et al*. 2017). We further characterize PREs in the context of inducible Cas9-directed genome editing. Despite seeing significant decreases in relative *Cas9* transcript levels, we are able to induce genome modifications with PREs upstream of the *Hsp70Bb* promoter when flies were reared at 18°C in the absence of a heat shock. This suggests that low levels of *Cas9* transcripts are sufficient to induce genome modifications and produce mutant mosaic phenotypes. Further modifications, either more PREs or in combination with temperature-sensitive inteins (Zeidler *et al*. 2004), are needed to completely suppress *Hsp70Bb* promoter activity with no heat shock to allow for precise control of expression of downstream genes.

### Many variables influence the effectiveness of PREs to reduce Cas9 efficacy with a heat-shock inducible promoter

Characterizing *PRE*-*Hsp70BbCas9* with gRNAs targeting three separate genes revealed that the gene target and/or the gRNAs themselves impact the downstream effect of a functional Cas9/gRNA complex, a common occurrence in studies of genome modification with Cas9. The very high rates (up to 100%) of biallelic mosaicism, by crossing Cas9 to *sgRNA:w* producing a clear visible phenotype, and lethal biallelic mosaicism by crossing to *dgRNA:Ser*, reveal this difference in comparison to *dgRNA:ey* which resulted in much lower rates of biallelic mosaicism in the form of a visible phenotype. Whether or not these differences in rate of visible biallelic mosaicism corresponds to different spatial expression levels of Cas9 mRNA and/or protein remains unknown. Effectiveness of PREs to reduce leaky expression of *Hsp70Bb* promoter is known to be influenced by genome integration site, proximal elements of the integration site, and tissue- and development-specific influences on Polycomb-based silencing (Dellino *et al*. 2004; Steffen and Ringrose 2014; Akmammedov *et al*. 2017). When generating *PRE-Hsp70BbCas9* systems, characterizing multiple transgene integration sites may aid in developing less-leaky inducible gene systems.

*D. melanogaster* rearing temperature appears to minimally affect levels of mutant formation by *Hsp70BbCas9* (without PREs) for efficacious gRNAs (*sgRNA:w* and *dgRNA:Ser*) at all heat shock durations. High rearing temperatures combined with long heat shocks appear to increase the rate of mutant phenotype for low efficient gRNA stocks (*dgRNA:ey*) without PREs, in our hands. Presence of PREs in *PRE-Hsp70BbCas9* allowed for more nuanced changes in rates of mutant phenotype formation for efficacious gRNAs (*sgRNA:w* and *dgRNA:Ser*). Higher rearing temperatures (21°C) for *PRE-Hsp70BbCas9* resulted in lower rates of *white* mutant phenotype formation compared to at 18°C after either no heat shock or 30 minute heat shocks (Figure 3). Often for *Serrate* mutant phenotype formation, higher rearing temperatures (26°C) with longer heat shock durations (1 hour or 2 hours) resulted in lower rates of visible biallelic mosaicism (Figure 5 H,J,M,N). Whether these numbers are a result of an increase in lethality remains unknown. Rearing temperature may be an important factor depending on the gene target and/or gRNA and details regarding whether or not these differences seen at varying rearing temperatures are due to differences in Cas9 function or PRE silencing ability remain for future studies.

### Using PREs in combination with genetic pest control strategies

A non-leaky, inducible promoter could be useful for generating gene drives (Champer *et al*. 2016) or other Cas9 modified fertile animals for large rearing and release while limiting the amount of Cas9 present in released animals. Lower levels of Cas9 RNA and protein may limit detrimental effects of Cas9 spreading through wild populations of pest species, depending on the gRNA and gene target, but this requires further exploration and detailed characterization for each gene drive system designed. Using CRISPR/Cas9 for sterile insect technique (SIT), as in precision-guided SIT (pgSIT), alleviates the concern for population spread of Cas9 since every released transgenic animal is sterile and therefore unable to pass on the transgene of concern (Kandul *et al*. 2019, 2021, 2022). A non-leaky temperature-inducible pgSIT (TI-pgSIT) would allow for chromosomal linkage of Cas9 and gRNAs without collapse of the transgenic stock population. This chromosomally linked TI-pgSIT stock would further eliminate the need to collect flies from separate strains; embryos could be heat shocked enmasse to generate lethal females and sterile males for release and allow for effective pest population control (Kandul *et al*. 2019, 2021).

## Supporting information

Reagents Table

## Data Availability

Strains are available through Bloomington *Drosophila* Stock Center (BDSC). Plasmids are available through Addgene. The Reagents Table contains details of all reagents used in this study.

## Acknowledgements

Thank you to Nikolay Kandul and Junru Liu for the use of the Hsp70Bb-Cas9-T2A-eGFP plasmid and *Hsp70BbCas9* transgenic *Drosophila melanogaster*. Thank you to Ting Yang for cloning advice.

## Author Contributions

O.S.A and N.W. conceived and designed the experiments. N.W., C.C., and C.R.D., performed molecular and genetic experiments. N.W. analyzed the data. All authors contributed to the writing and approved the final manuscript.

## Ethical conduct of research

All samples were handled in accordance with the UCSD Biological Use Authorization (BUA #R2401).

## Supplemental Materials

**Supplemental Figure S1.**
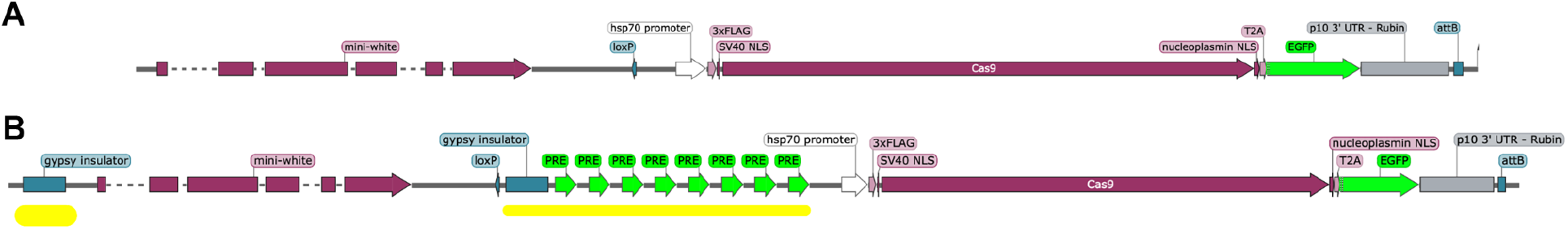
Transgene maps to compare *Hsp70BbCas9* and *PRE-Hsp70BbCas9*. A) Hsp70Bb-Cas9-T2A-eGFP (part of Addgene 153284) used to generate *Hsp70BbCas9* transgenic *D. melanogaster*. B) PRE-Hsp70Bb-Cas9_1.3 (part of Addgene 190797) used to generate *PRE-Hsp70BbCas9* transgenic *D. melanogaster*. Differences between the transgenes are highlighted in yellow and consist of the Polycomb response elements (PREs) (neon green) and gypsy insulator elements (teal). Both transgenes were inserted using ϕC31-mediated integration into the same genomic location (see Materials and Methods).

**Supplemental Figure S2.**
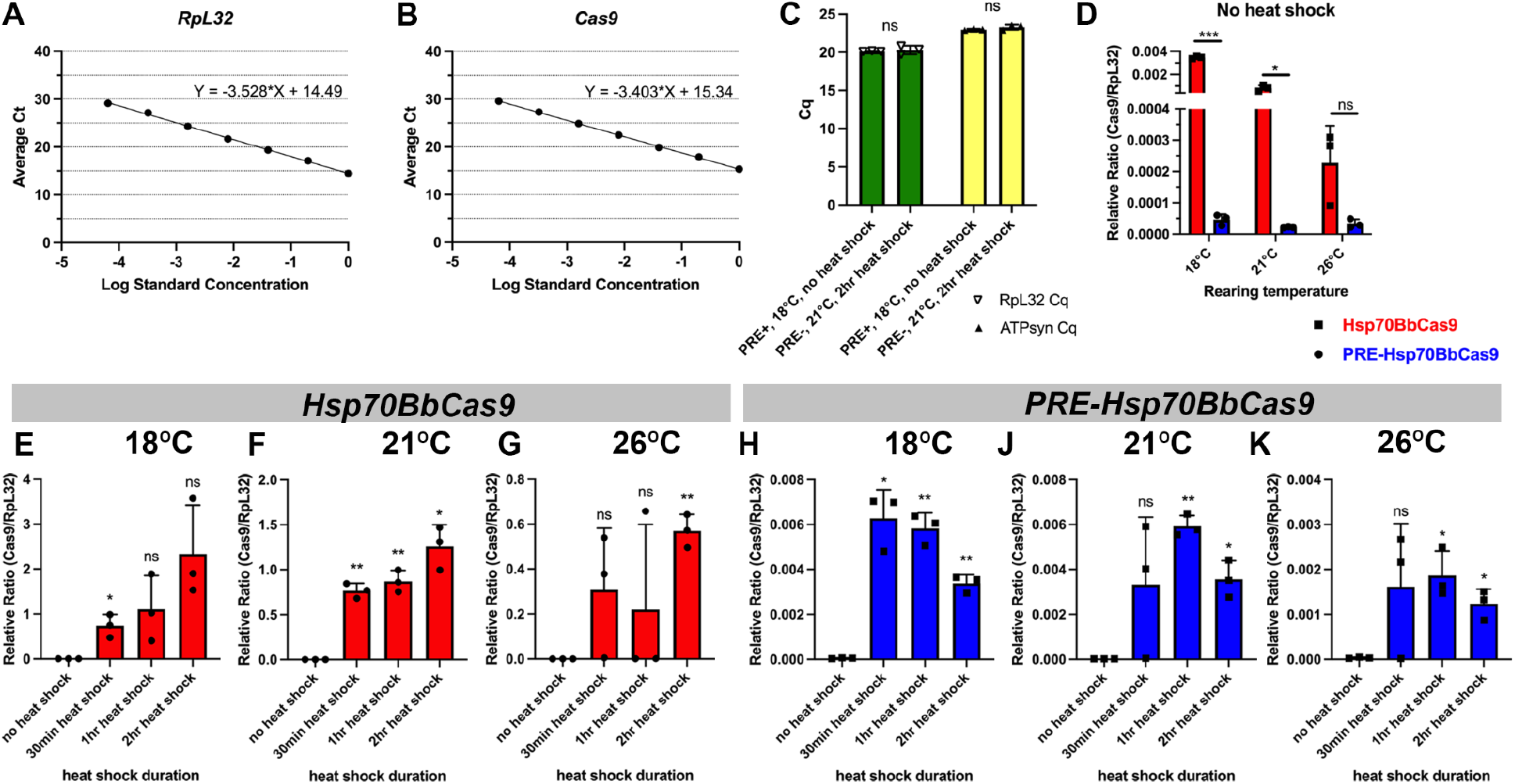
Effects of Polycomb response elements on relative *Cas9* transcript levels. Standardization of A) *RpL32* primers and B) *Cas9* primers. C) Comparison of Cq values for house-keeping genes *RpL32* (green) and *ATPsyn* (yellow). D) Relative ratio (E _*RpL32*_ ^RpL32_Ct^/E_*Cas9*_ ^Cas9_Ct^) of *Cas9*/*RpL32* transcripts after no heat shock at varying rearing temperatures, with or without PREs. Relative ratio (E _*RpL32*_ ^RpL32_Ct^/E *Cas9* ^Cas9_Ct^) of *Cas9*/*RpL32* transcripts at varying rearing temperatures and heat shock durations for E-G) *Hsp70BbCas9* and H-K) *PRE-Hsp70BbCas9* (no heat shock data reanalyzed from C). Hsp (red) = *Hsp70BbCas9*, PRE (blue) = *PRE-Hsp70BbCas9*. Error bars = standard deviation. Significance calculated using unpaired two-tailed t-test with Welch’s correction. ns = P > 0.05, * = P ≤ 0.05 ** = P ≤ 0.01, *** = P ≤ 0.001, **** = P ≤ 0.0001. Statistics for E-K were calculated for each heat shock duration (30 minute, 1 hour, 2 hour) compared to no heat shock. Please note differences in y-axis scaling.

**Supplemental Figure S3.**
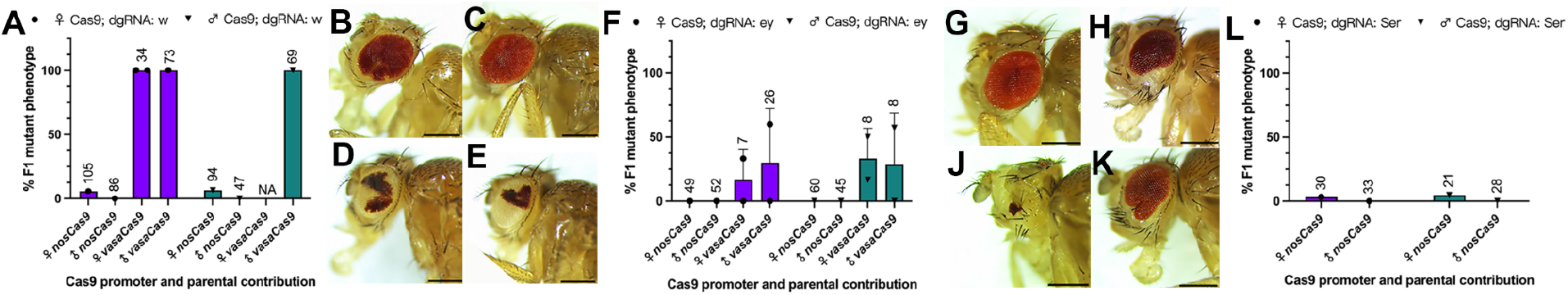
Mutant phenotype formation induced by either *nanosCas9* or *vasaCas9* controls. Quantification of F1 mutant phenotypes for A) *sgRNA:w*, F) *dgRNA: ey*, and L) *dgRNA:Ser* with either *nosCas9* or *vasaCas9* inherited maternally or paternally. *vasaCas9/dgRNA:Ser* trans-heterozygous flies (maternal and paternal Cas9) all died during pupal development and were not scored/reported in this figure. Purple = female trans-heterozygous, green = male trans-heterozygous. n = number of flies scored (listed above corresponding bar). Error bars = standard deviation. Significance was not calculated. B-E) Representative images of the phenotypic range of Cas9-induced *white* phenotypes with B) maternal *nosCas9*, C) paternal *nosCas9*, D) maternal *vasaCas9*, E) paternal *vasaCas9*. G-K) Representative images of the phenotypic range of Cas9-induced *eyeless* phenotypes with G) paternal *nosCas9*, H) maternal *vasaCas9*, J-K) paternal *vasaCas9*. All scale bars = 250μm.

**Supplemental Figure S4.**
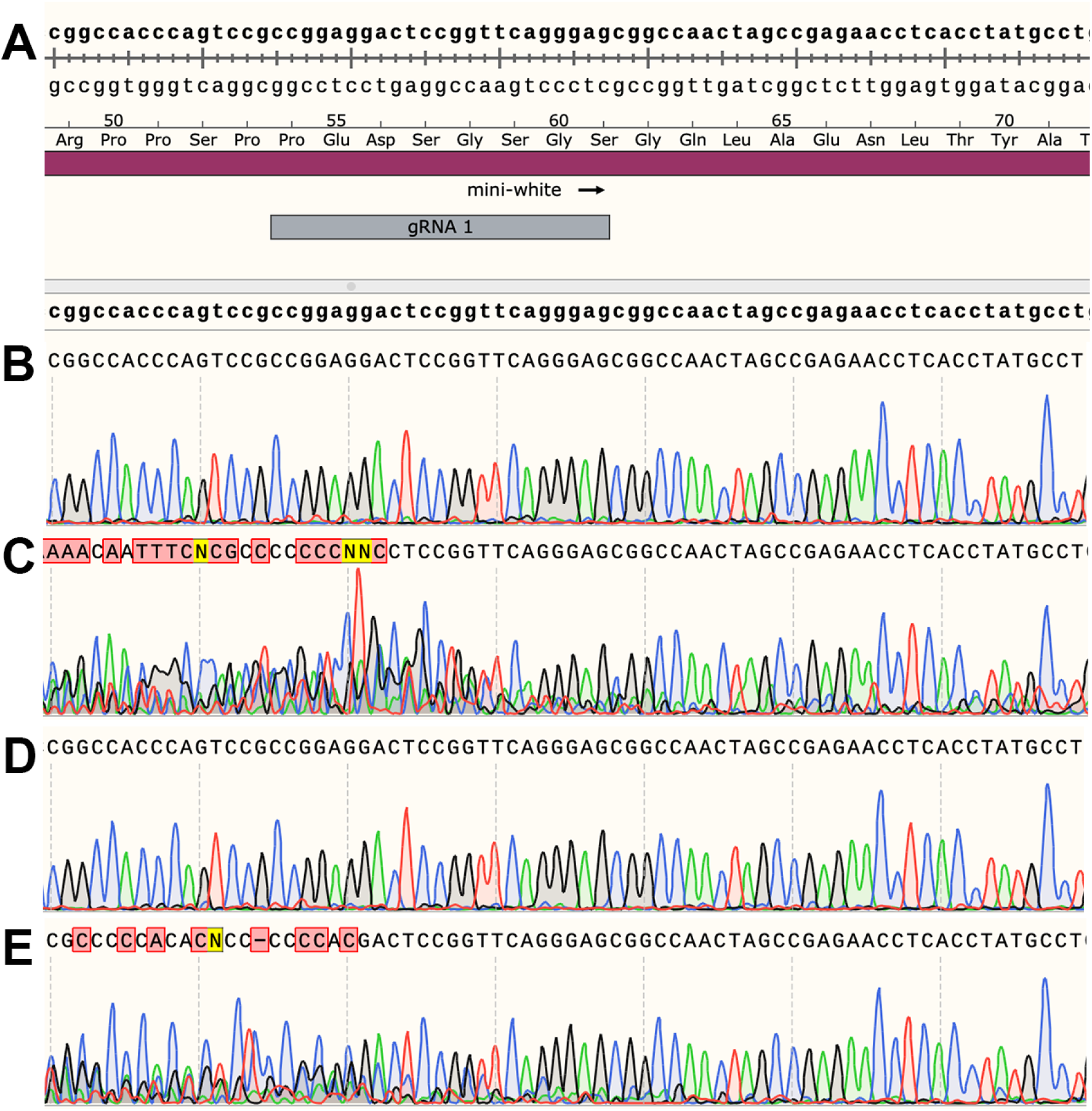
Sanger sequencing genotyping of heat-shock-induced Cas9 targeting of *mini-white*. F1 generated from *Hsp70BbCas9* crossed with *sgRNA:w* were genotyped for mutations at the *mini-white* transgene since the F1 had heterogenous *white* alleles which Sanger sequencing cannot distinguish. F1 from *PRE-Hsp70BbCas9* crosses were not genotyped. A) Transgene sequence from plasmid Hsp70Bb-Cas9-T2A-eGFP, B) maternal *Hsp70BbCas9* heterozygous control, C) maternal *Hsp70BbCas9* trans-heterozygous with a phenotype, D) paternal *Hsp70BbCas9* heterozygous control, E) paternal *Hsp70BbCas9* trans-heterozygous with a phenotype.

**Supplemental Figure S5.**
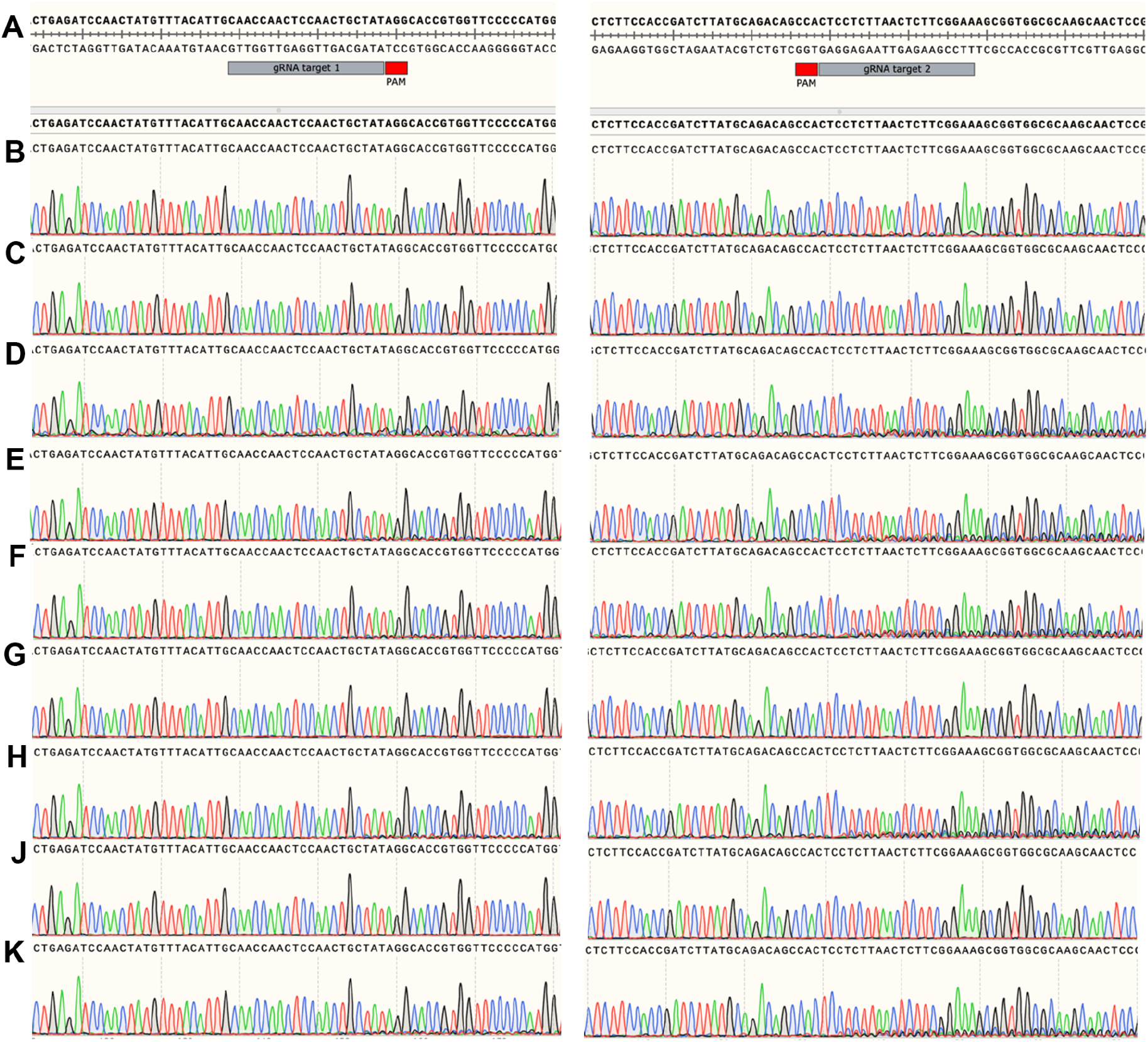
Sanger sequencing genotyping of heat-shock-induced Cas9 targeting of *eyeless*. Left column = chromatograms of gRNA target 1 sequence. Right column = chromatograms of gRNA target 2 sequence. A) Reference *D. melanogaster* genome sequence of *eyeless* from NCBI Reference Sequence: NC_004353.4, B) *w*^*1118*^, C) maternal *Hsp70BbCas9* heterozygous control, D) maternal *Hsp70BbCas9* trans-heterozygous with a phenotype, E) paternal *Hsp70BbCas9* heterozygous control, F) paternal *Hsp70BbCas9* trans-heterozygous with a phenotype, G) maternal *PRE-Hsp70BbCas9* heterozygous control, H) maternal *PRE-Hsp70BbCas9* trans-heterozygous without a phenotype, J) paternal *PRE-Hsp70BbCas9* heterozygous control, K) paternal *PRE-Hsp70BbCas9* trans-heterozygous with a phenotype.

**Supplemental Figure S6.**
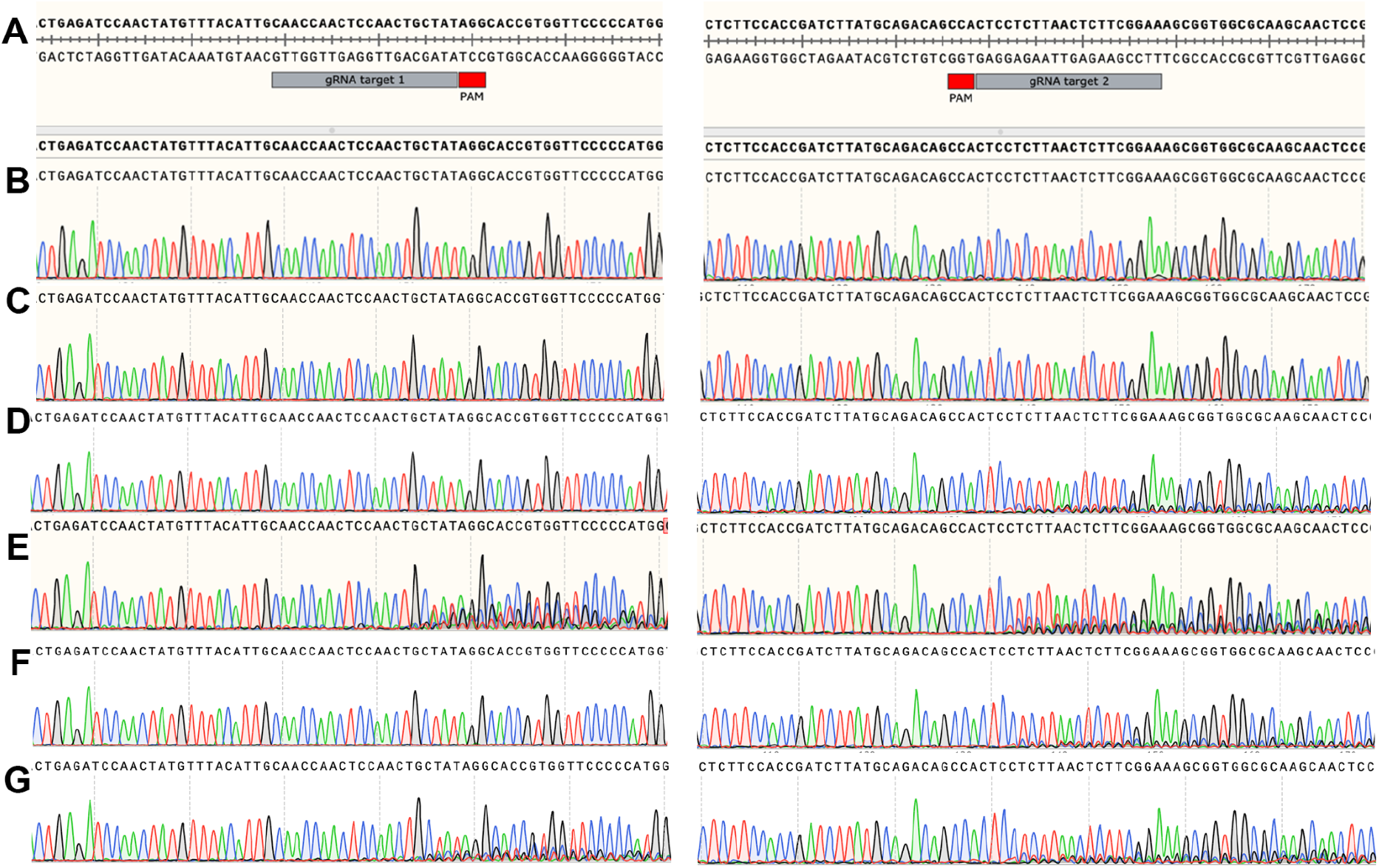
Sanger sequencing genotyping of *nanos* or *vasa* promoter-driven Cas9 targeting of *eyeless*. Left column = chromatograms of gRNA target 1 sequence. Right column = chromatograms of gRNA target 2 sequence. A) Reference *D. melanogaster* genome sequence of *eyeless* from NCBI Reference Sequence: NC_004353.4, B) *w*^*1118*^, C) F1 from ♀*nosCas9* x ♂*dgRNA:ey* without a phenotype, D) F1 from ♀*dgRNA:ey* x ♂*nosCas9* without a phenotype, E) F1 from ♀*vasaCas9* x ♂*dgRNA:ey* with a phenotype, F) F1 heterozygous control from ♀*dgRNA:ey* x *vasaCas9*, G) F1 from ♀*dgRNA:ey* x *vasaCas9* with a phenotype.

**Supplemental Figure S7.**
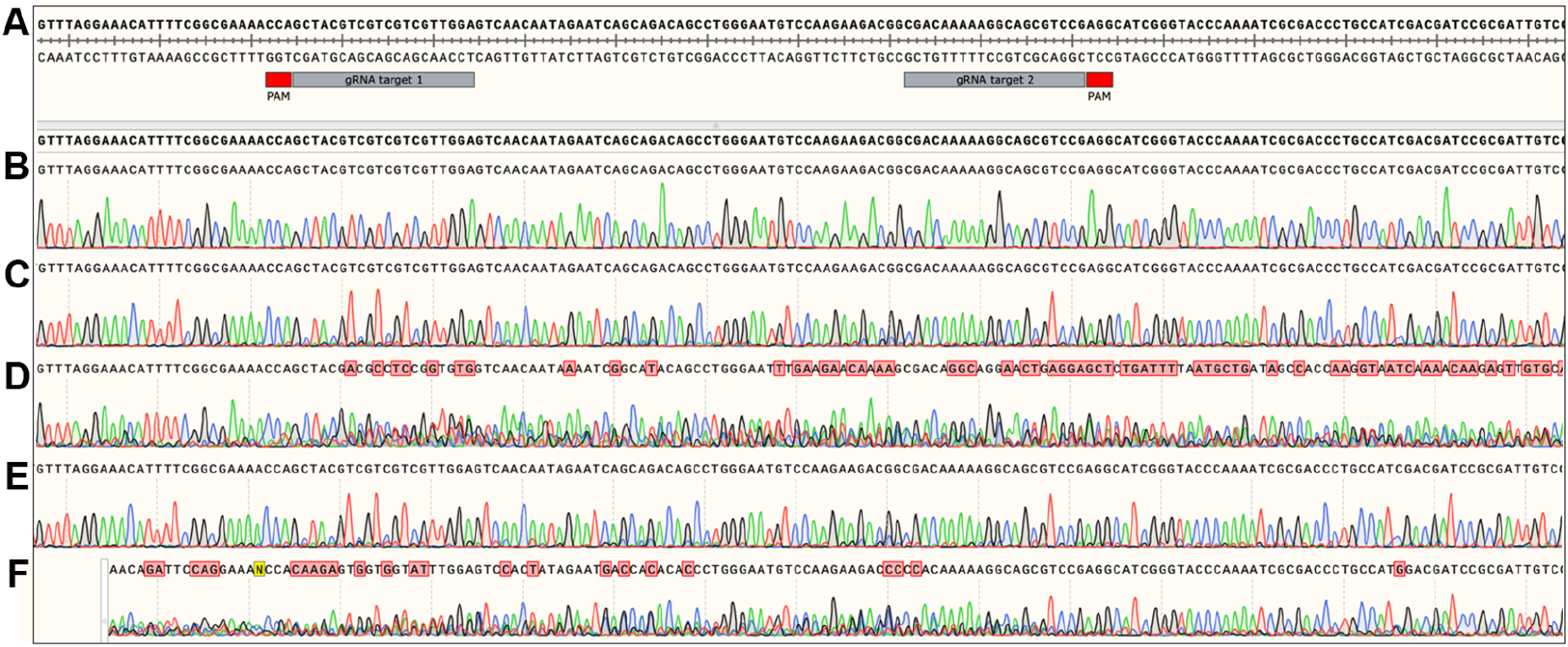
Sanger sequencing of heat-shock induced *Hsp70BbCas9* targeting of *Serrate*. A) Reference *D. melanogaster* genome sequence of *Serrate* from NCBI Reference Sequence: NT_033777.3. gRNA target sequences are annotated in gray boxes, PAMs annotated in red boxes. Sequencing calls and corresponding chromatograms for B) *w*^*1118*^, C) maternal *Hsp70BbCas9* heterozygous control, D) lethal pupae with maternal *Hsp70BbCas9*, E) paternal *Hsp70BbCas9* heterozygous control, F) lethal pupae with paternal *Hsp70BbCas9*.

**Supplemental Figure S8.**
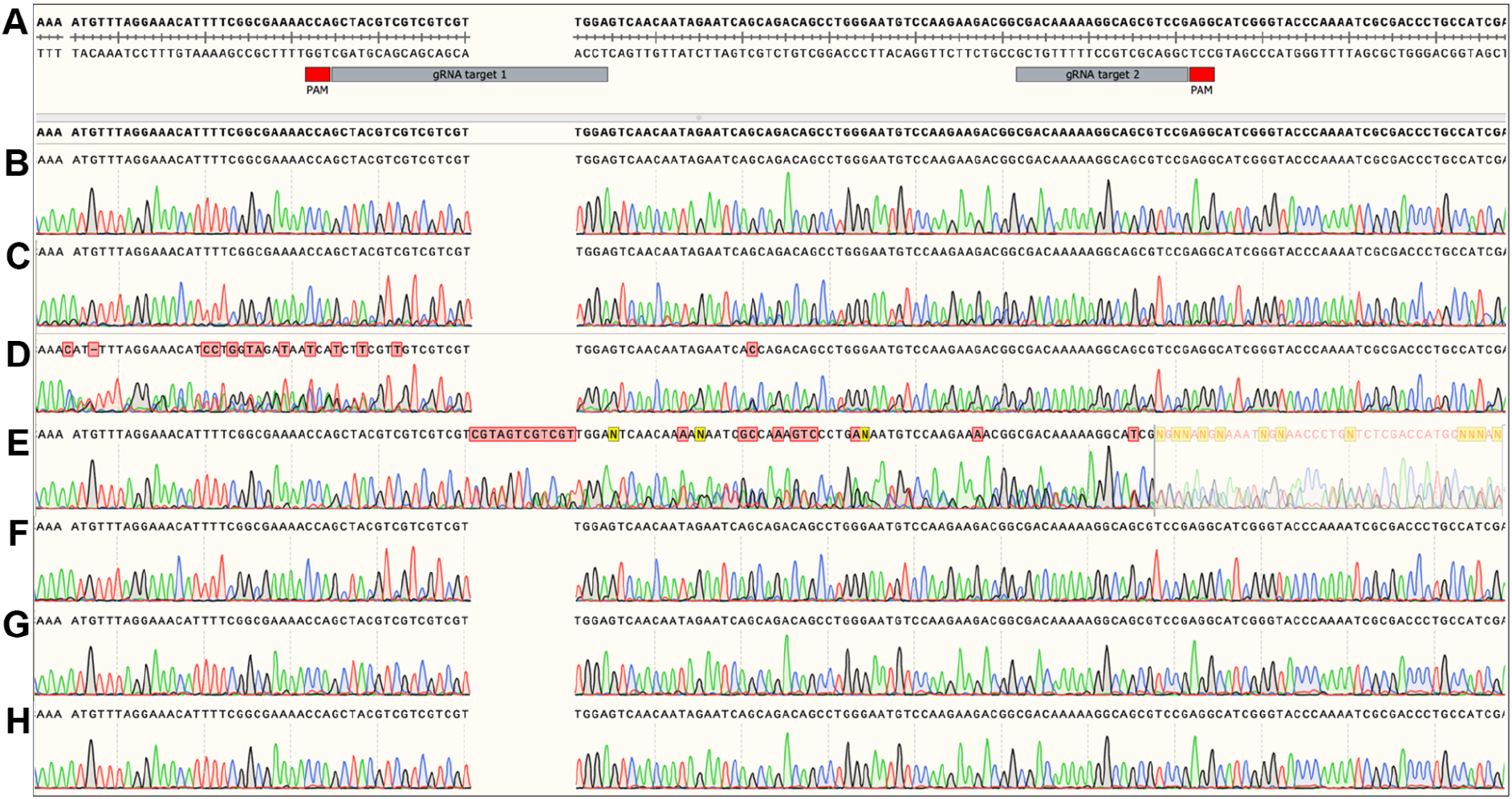
Sanger sequencing of heat-shock induced *PRE*-*Hsp70BbCas9* targeting of *Serrate*. A) Reference *D. melanogaster* genome sequence of *Serrate* from NCBI Reference Sequence: NT_033777.3. gRNA target sequences are annotated in gray boxes, PAMs annotated in red boxes. Sequencing calls and corresponding chromatograms for B) *w*^*1118*^, C) maternal *PRE-Hsp70BbCas9* heterozygous control, D) maternal *PRE-Hsp70BbCas9* trans-heterozygote without serrated wing phenotype, E) maternal *PRE-Hsp70BbCas9* trans-heterozygote with serrated wing phenotype, F) paternal *PRE-Hsp70BbCas9* heterozygous control, G) paternal *PRE-Hsp70BbCas9* trans-heterozygote without serrated wing phenotype, H) paternal *PRE-Hsp70BbCas9* trans-heterozygote with serrated wing phenotype.

**Supplemental Figure S9.**
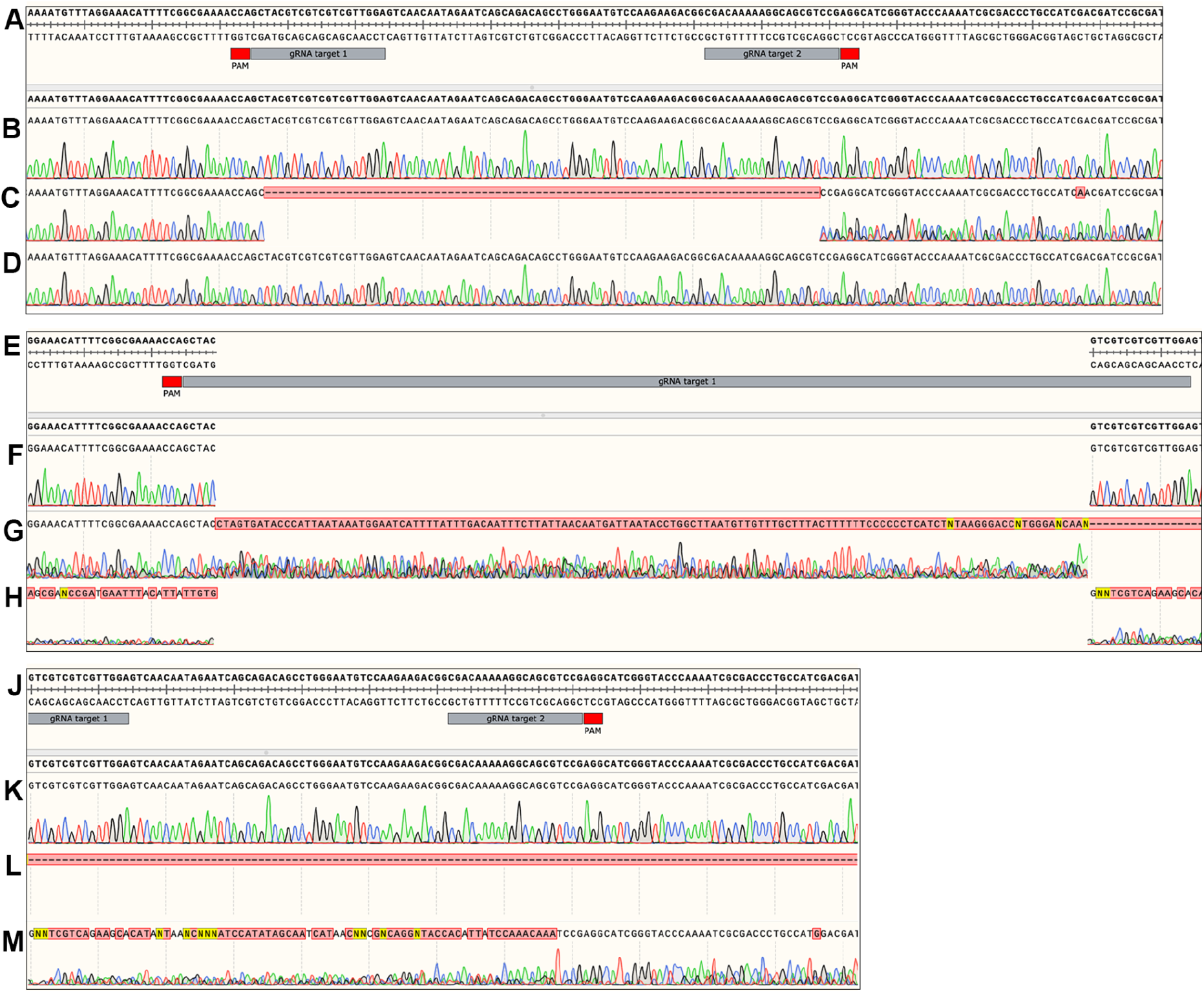
Sanger sequencing genotyping of *nanos* or *vasa* promoter-driven Cas9 targeting of *Serrate*. A) Reference *Drosophila melanogaster* genome sequence of *Serrate* from NCBI Reference Sequence: NT_033777.3. gRNA target sequences are annotated in gray boxes, PAMs annotated in red boxes. Sequencing calls and corresponding chromatograms for B, F, J) *w*^*1118*^, C) F1 from ♀*nosCas9* x ♂*dgRNA:Ser*, D) F1 from ♀*dgRNA:Ser* x ♂*nosCas9*, E) reference (as in A) for gRNA1, F) *w*^*1118*^, G) F1 ♀*vasaCas9* x ♂*dgRNA:Ser*, gRNA1, H) ♀*dgRNA:Ser* x *vasaCas9*, gRNA1, J) reference (as in A) for gRNA2, K) *w*^*1118*^, L) ♀*vasaCas9* x ♂*dgRNA:Ser*, gRNA2, M) ♀*dgRNA:Ser* x *vasaCas9*, gRNA2.

